# Active sampling in visual search is coupled to the cardiac cycle

**DOI:** 10.1101/405902

**Authors:** Alejandro Galvez-Pol, Ruth McConnell, James M. Kilner

## Abstract

Recent research has demonstrated that perception and reasoning vary according to the phase of internal bodily signals such as heartbeat. This has been shown by locking the presentation of sensory events to distinct phases of the cardiac cycle. However, task-relevant information is not usually encountered in such a phase-locked manner nor passively accessed, but rather actively sampled at one’s own pace. Moreover, if the phase of the cardiac cycle is an important modulator of perception and cognition, as previously proposed, then the way in which we actively sample the world should be similarly modulated by the phase of the cardiac cycle. Here we tested this by coregistration of eye movements and heartbeat signals while participants freely compared differences between two visual arrays. Across three different analyses, we found a significant coupling of saccades, subsequent fixations, and blinks with the cardiac cycle. More eye movements were generated during the systolic phase of the cardiac cycle, which has been reported as the period of maximal effect of the baroreceptors’ activity upon cognition. Conversely, more fixations were found during the diastole phase (quiescent baroreceptors). Lastly, more blinks were generated in the later period of the cardiac cycle. These results suggest that interoceptive and exteroceptive processing do adjust to each other; in our case, by sampling the outer environment during quiescent periods of the inner organism.

## 1. Introduction

Interoception refers to the set of physiological and cognitive processes that are involved in determining the physiological condition of the body (Craig, 2002). It can be distinguished from the domains of exteroception (processing of the environment) and proprioception (the position of the body in space) as a distinct sensory domain. Recently, research studying interoception has markedly increased (Khalsa et al., 2018; Khalsa & Lapidus, 2016). There are at least two motivations for this renewed interest: establishing the extent to which interoception contributes to cognition in the neurotypical brain and assessing the clinical impact of atypical interoception.

The primary physiological focus of most studies on interoception is the cardiovascular system. In a single heartbeat/cardiac cycle two main phases are observed. In the systolic phase the heart contracts and ejects the blood, whereas in the diastole phase the heart expands while being filled. Both phases comprise a whole heartbeat, with the R-peak (peak in electrocardiogram depicting the contraction at systole) indicating the start of a new heartbeat. Since early evidence of a modulatory effect of the carotid sinus in both autonomous and central nervous system (Koch, 1932; Kreindler, 1946), research in humans have shown that participants’ response to stimuli changes according to the phase of the heartbeat in which the information is presented (Azevedo, Badoud, & Tsakiris, 2017; Critchley & Garfinkel, 2018; Salomon et al., 2016). For instance, in the sensorimotor domain, processing information during systole has been associated to the reduction of sensitivity towards pain, visual, and auditory stimuli (Edwards et al., 2001; Edwards et al., 2007; McIntyre et al., 2006; Pramme et al., 2014). Cardiac afferent signals seem to play an important role in the interaction of the heart-brain axis, moderating cognitive processes, competing for the allocation of attentional and representational resources, and dampening/enhancing sensory processing (Critchley & Garfinkel, 2018; Critchley & Harrison, 2013; Khalsa et al., 2018; Seth, 2013).

The demonstrations that perception and cognition are modulated as a function of the cardiac cycle have been achieved by deliberately time-locking the presentation of stimuli to the participants during diastole or systole phases. However, in our everyday experiences sensory information is not presented phase-locked to the cardiac cycle. Therefore, although of considerable interest, it is unclear how these findings impact on our cognition outside of the laboratory setting. If the phase of the cardiac cycle is an important modulator of perception and cognition, as previously proposed, then the way in which we freely and actively sample the world should be similarly modulated by the phase of the cardiac cycle. The current study was designed to test this hypothesis. To our knowledge only two studies have recently started to examine this. Ohl and colleagues (2016) reported the generation of involuntary phase-locked microsaccades to the systolic phase of the cardiac cycle. Similarly, Kunzendorf et al., (2019) reported the generation of more keypresses, which led to the onset of images to-be-sampled, during the systolic phase. Here we tested the hypothesis that the timing of eye movements in a free visual search task would be modulated during the different phases of the cardiac cycle. We hypothesized that the generation of more phase-locked eye movements (saccades) during systole and the opposite pattern (absence of eye movements; fixations) during diastole; i.e., we tested whether or not the timing of eye movements in a free visual search task would be modulated by the phases of the cardiac cycle. To this end, we coregistered oculomotor behaviour and electrocardiogram (ECG) while human participants performed a visual search task comparable to ‘spot the difference’. Specifically, we recorded the position of the eyes to detect saccades, fixations, and blinks while participants compared differences in colouration between two bilateral arrays (Figure 1a). For each heartbeat, we computed the point at the phase of the cardiac cycle in which each oculomotor event occurred. The prediction was that if participants’ behaviour is modulated by their cardiac signals, the eye movements generated during the present free visual task would reflect this (more saccades, fixations, and blinks occurring in particular phases of the cardiac cycle).

**Figure 1.**
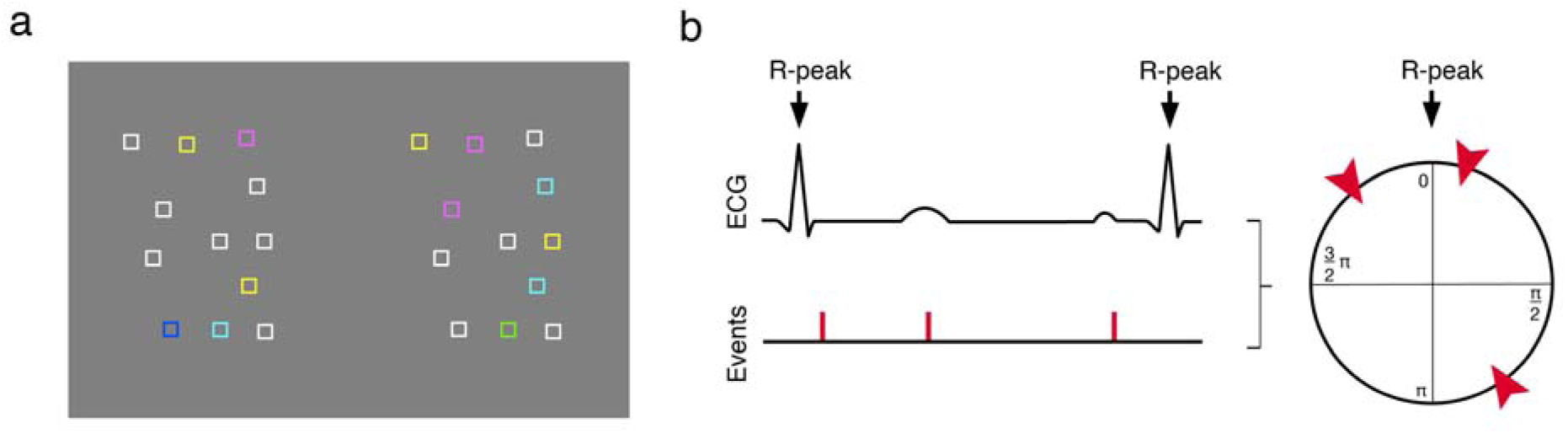
Task and schematic illustration of cardiac and oculomotor events. (a) Shows an example of a single trial. Participants were instructed to report the total number of boxes that differed in colour between two bilateral arrays. For this, they compared each box in the left array with the homologous-positioned box in the right array. If the colour was different, it counted as a difference; the trial shown above depicts 9 differences. (b) A schematic of the analysis. For instance, for each saccade onset event we computed its phase (in degrees) relative to the concurrent heartbeat: for an R-R interval of time t_R_, where the saccade onset occurred at time t_E_, we calculated t_E_/T_R_ x 360. For each participant, the mean phase was then calculated separately for the saccades and blinks onset, as well as for the mean point of fixations by using circular averaging

## 2. Materials and methods

### 2.1. Participants

32 healthy adults participated (24 females; age range = 18-39). All participants reported both normal or corrected-to-normal vision and normal cardiac condition. All subjects gave informed consent, were reimbursed for the participation, and volunteer to take part in the experiment. Ethical approval for the methods/procedures was obtained from the UCL research ethics committee.

### 2.2 Task and procedure

Participants received instructions and completed two practice trials. Next, they completed 5 blocks of 20 trials while their electrocardiogram (ECG) and oculomotor behaviour were recorded. Participants performed a visual search comparable to a controlled version of ‘the spot the difference task’. Participants were instructed to count the total number of boxes distinctively coloured across two bilateral arrays that were continuously displayed on the screen. A given difference consisted in dissimilar colour between a box in one array and the homologous-positioned box of the other array (Figure 1a). Participants were explicitly told that they could take as long they felt in each trial and they advanced to the next trial by pressing the space-bar and reporting the number of differences. In addition to the coloured boxes, catch trials (changing one of the boxes by a small red circle) were included in 3% of the trials to enforce ongoing attention to the task. Participants had to report the sight of these latter stimuli and count their presence as a difference. All participants reported the presence of these red circles in the catch trials.

### 2.3 Visual display

The visual arrays were displayed on a 17-inches LCD monitor (60 Hz) and resolution of 1280 x 800 pixels. Participants were seated in a dimly light room at 66 cm from the monitor with their head on a chin rest and their forearms rested on the top of a table. The visual arrays were generated and displayed using Matlab R2016b (The MathWorks Inc., Natick, MA, USA) with the Psychtoolbox-3 (Brainard, 1997; Cornelissen, Peters, & Palmer, 2002; Kleiner et al., 2007; Pelli, 1997). We recorded oculomotor behaviour while participants compared differences in coloration between small coloured boxes (0.8° x 0.8° each) displayed within two alike bilateral arrays embedded in two rectangular regions (8° x 11°) centred 6° to the left and right side of the screen centre. The position of the boxes was randomised with the restriction that the distance between them greater than 2.2° (centre to centre). To decide the assignment of colouration in each frame, we previously run several pilots in which we also considered the task’s length and difficulty, and the number of generated saccades and heartbeats. In each trial, each coloured frame was subjected to a combination of the RGB triplet, assigning a .18 probability of obtaining 1 in each individual number of the triplet (see https://bit.ly/2jUvdFd). Overall, this equals to a ~.58 likelihood of obtaining a white frame. In the remaining probability a different colour out of the remaining seven RGB combinations was obtained (approximately 8 differences in average).

### 2.4. Eye tracker recording and pre-processing

Eye positions were recorded from the left eye using a video-based eye tracking System (Eyelink 1000 Plus; SR Research, Ottawa, ON, Canada); sampling rate 1000Hz. Before each block, a standard nine points calibration procedure controlled the correspondence between the position of the pupil and the screen display. Also, before each trial a single centred target was presented to allow estimating the drift between the current fixation and the above-mentioned calibration. This procedure was launched via EyelinkToolbox through Psychtoolbox-3 and run in Matlab R2016b. At the onset of each trial, a squared pulse was delivered through a data acquisition box (NI USB-6229; National Instruments Corp., ATX, USA) to a CED Power unit 1401-3A, which signal was recorded in simultaneity to the ECG in Spike2 8.10 (Cambridge Electronic Design Limited, Cambridge, UK). This pulse allowed us to align the oculomotor and ECG recordings.

In order to detect oculomotor events, we used the parsing system for cognitive research implemented in the eye tracker. A saccade event was detected if the velocity and acceleration of an eye movement reached specified thresholds; these involve motion, velocity, and acceleration (0.1°, 30°/sec, and 8000°/sec^2^, respectively). Despite the task was designed to elicit voluntary, large, and clear saccades, a small portion of microsaccades (<1.5 °) was also generated/recorded. Heartbeats with these latter pattern of eye movements were excluded from further analyses.

### 2.5. ECG recording and pre-processing

ECG was recorded using a D360 8-Channel amplifier (Digitimer Ltd, Hertfordshire, UK) in Spike2 8.10; sampling rate 1000 Hz. The electrodes were placed over the right clavicle and left iliac crest according to Einthoven’s triangular arrangement. To identify each heartbeat, we detected the R peaks in the ECG trace by computing vectors with the local maxima of the ECG; minimum peak to peak distance of ~500 ms and a mean minimum peak height of ~0.7 millivolts. We visually inspected the detection of the R-peaks returned by the algorithm and applied small adjustments when required. This visual inspection also allowed us to observe small drifts impairing the correct detection of all the R-peaks, due for instance, to some participants producing small body movements. Given the temporal consistency between R-peaks, oculomotor events occurring in R-R intervals separated by a distance ±1.25 x the median of the participants’ R-R interval in milliseconds were not considered in further analyses.

## 3. Data analysis

To examine whether or not there was a significant statistical relationship between the saccades, fixations, blinks, and the phase of the cardiac cycle, we calculated the phase of these oculomotor events as a function of the R-R interval (Kunzendorf et al., 2019). Circular statistics were employed in order to exploit the iterating nature of the cardiac cycle. For each oculomotor event, the time of the preceding and subsequent R-peak were calculated. Then, the phase of the generated oculomotor event was computed as a function of the R-R interval in which it occurred (Figure 1b). This results in three mean phases per participant, one for each type of oculomotor event. Next, we tested separately for each condition whether these phases differed from a uniform distribution (‘dispersed’ non-phased oculomotor events) using Rayleigh tests. We averaged these distributions across participants for saccades onset, mean time point of fixations, and blinks onset, indicating the degree of uniformity of the phases for each participant.

## 4. Results

### 4.1. Descriptive results

The mean number of computed heartbeats across the experiment was 1211, SD = ± 315.3. The mean heart rate of the participants was 72 bpm; 834ms from R to R peak, SD = ± 145ms. The removal of oculomotor events in noisy sections of the ECG (see *Methods*) comprised 7% of the total number of events. After removal of noisy sections of the ECG, the average number of saccades and fixations computed was 3109.8, SD = ± 750. The average number of blinks was 109.3, SD= ± 118.1. The average duration of the saccades was 47 ms, SD = ±5.2 ms; their average amplitude was 8°, SD = ±0.8. The average duration of the subsequent fixations was 212.7 ms, SD = ±26 ms. Participants generated 3.7 saccades per second (Hz), SD = ±0.34. The average number of blinks was 109.3, SD = ± 118.1, and their average duration was 230ms, SD = ±57.4 ms. The completion of each trial during the task took approximately 8.3 sec.

### 4.2. Statistical results

We proceed to test our main hypothesis, the coupling of the participants’ gaze (saccades, fixations, and blinks) with their own heartbeat. The dataset used for these analyses is publicly available via Open Science Framework in the following link: https://osf.io/h5x84/. We first analyzed the saccades onset, which occurred on average between heartbeats at 101.7 ° (Figure 2a). This distribution differed significantly from the uniform distribution (Rayleigh test z = 5.220, p = 0.0047) and it was positively skewed around ~100 degrees (~1/2π). The mean phase where statistically more saccades were generated corresponds to the early period of the cardiac cycle, systolic phase. This period has been linked to maximal baroreceptor influence upon cognition (Azevedo et al., 2017; Edwards et al., 2001; Garfinkel et al., 2014; Garfinkel & Critchley, 2016; McIntyre et al., 2006). Secondly, we also computed the mean time point of fixations for each fixation by using circular averaging. On average participants fixated between heartbeats at 206.2 ° (Figure 2b). This distribution differed significantly from the uniform distribution (Rayleigh test z = 9.534, p = 0.000037) and it was positively skewed around ~206 degrees. The mean phase where statistically more fixations were generated would correspond to the mid phase of the cardiac cycle (early diastole), which has been reported as the quiescent period of baroreceptor activity (Critchley & Harrison, 2013). Last, we separately computed those heartbeats with concurrent blinks, which were not included in the above analyses. The rationale behind this is that upon examination of blinks, these data seem qualitatively different. Blinks accounted for a smaller number of events when compared to saccades and fixations. Moreover, they might affect the ‘two-sided’ pattern of eye gaze found in the vast majority of heartbeats, which comprise eye movements and the absence of these (fixations). Blinks onset occurred on average at 269.0 ° (Figure 2c). This distribution differed significantly from the uniform distribution (Rayleigh test z = 7.6, p = 0.00034) and it was positively skewed around ~270 degrees (~3/2π). The mean phase where statistically more blinks were generated would correspond to the late period of the cardiac cycle, during the diastole phase.

**Figure 2.**
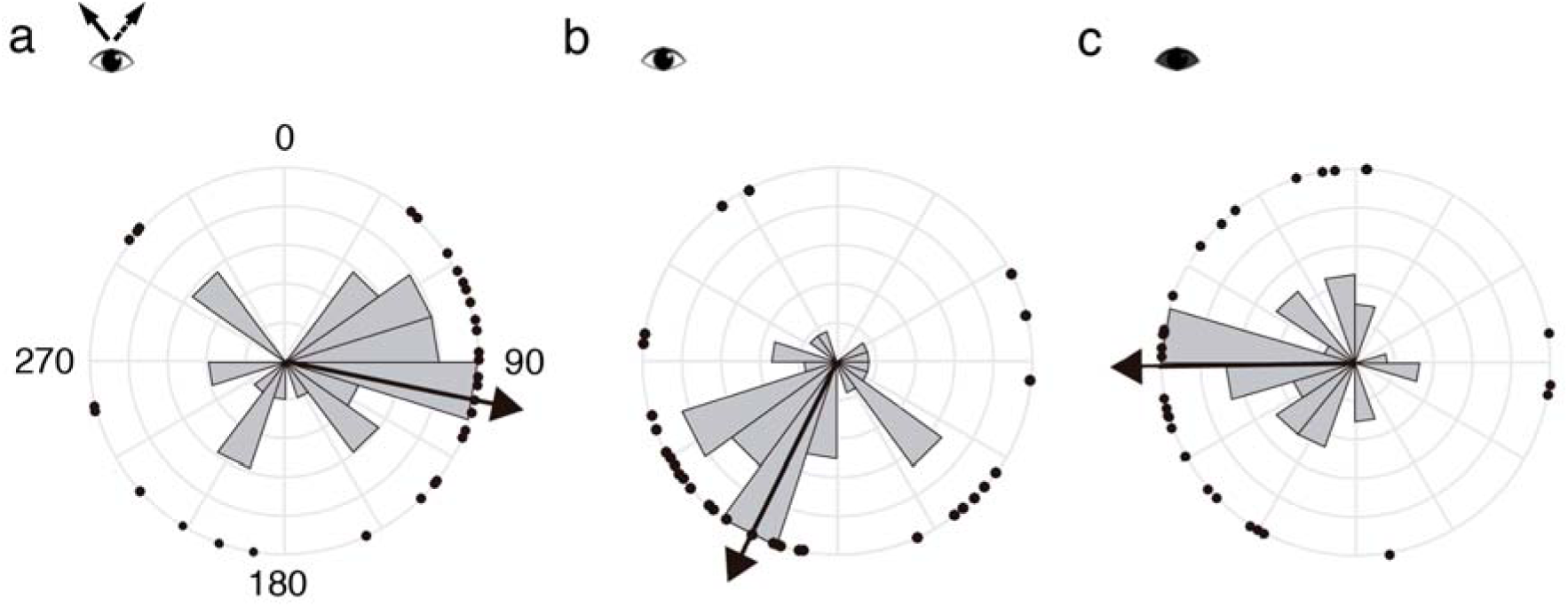
Relationship between patterns of eye gaze and cardiac cycle. In each panel the R-peaks occurred at 0 ° and the phase through the R-R interval cycles clockwise round the circle. Panel (a) shows the relationship between saccades onset with respect of the phase of the R-R interval. (b) The figure shows the relationship between fixations mean phase with respect to the R-R interval. (c) The figure shows the relationship between blinks onset with respect of the phase of the R-R interval. The black dots show the mean phase for each participant. The circular histogram shows the frequency of data as a function of phase; max. histograms equals to 6 participants. The solid black arrow shows the circular mean phase across subjects. Saccades onset, mean time point of fixations, and blinks onset occurred significantly more often in the early, mid, and later period of the cardiac cycle, respectively.

In summary, the results of the circular statistics show that more saccades were generated during the early phase of the cardiac cycle (systole) whereas for fixation and blinks more fixations were generated during the mid and late phase of the cardiac cycle (diastole). In addition to the current analyses, we also computed the change in proportion of saccades, fixations, and blinks along the cardiac cycle in 6 bins that were equally spaced throughout the R-R interval (analysed using a repeated-measures ANOVA with one factor with 6 levels), as well as the inter-trial-phase-coherence, whereby a vector of a certain length and direction represents the strength of the observed effects. The analyses, results, and subsequent figures are depicted in supplementary materials. Overall, these yielded similar results to those from the circular statistics (i.e., significant modulation of saccades, fixations, and blinks along the cardiac cycle).

## 5. Discussion

The effect of the cardiac cycle upon cognition has been observed by meticulously time-locking the presentation of information to each cardiac phase (Azevedo et al., 2017; Garfinkel et al., 2014; Park et al., 2017). Here we foster a more ecological framework in which participants may ‘take in’ information at their own pace according to cardiac signalling. To examine this, we allowed participants to freely sample task-relevant visual information while recording their oculomotor behaviour and ECG. Our analyses revealed a significant coupling of saccades, subsequent fixations, and blinks to different periods along the cardiac cycle. The mean phase of saccades onset, mean time point of fixations, and blinks onset occurred at 101, 206, and 269 degrees (i.e., ~1/4, 2/4, and 3/4 of a complete 360 ° cycle, respectively). According to Mao et al., (2003), who examined the percentage of the whole cardiac cycle comprising the systolic and diastole phases in more than 800 people, our above-mentioned mean phases occurred in the mid systole, early diastole, and mid diastole (e.g., systole takes approximately the first 40% of the whole cycle, systole the remaining).

In primates, only the central portion of our visual system has sufficient receptors to obtain visual information with high resolution. Therefore, we generate eye movements to sample elements of interest distributed in the environment. This sampling encompasses changes in object representation, framing, allocation of attentional resources (Hamker et al., 2008; Ibbotson & Krekelberg, 2011), as well as allows collecting sensory data to test hypotheses and beliefs about the world (see eg., Friston et al., 2012; Parr and Friston, 2018). Here combining perceptual and motor mechanisms allows the reduction of entropy (Friston et al., 2012, 2009; Pickering & Clark, 2014). Interestingly, visual perception is impaired during saccades. Previous research has described the selective suppression of the magnocellular pathway during saccades (Burr, Morrone, & Ross, 1994; Ross, Burr, & Morrone, 1996). Moreover, one the processes involved in the current task (i.e., serial search and scanning) seem to involve this pathway (Cheng, Eysel, & Vidyasagar, 2004). Conversely, other studies have described that during the postsaccadic period, visual acuity is enhanced (as reviewed by Ibbotson and Krekelberg, 2011).

Our results show that participants generated more saccades during the early phase of the cardiac cycle (systole) than in the later diastole. The systole period is concurrent with sensors on vessels, namely baroreceptors, that fire during the heart’s ejection of blood. The fixations generated during the task follow the opposite pattern, the mean phase of the fixations was found consistently during diastole (quiescent period of the baroreceptors). Thus, one possibility to be further examined is that participants partly generated more saccades, in which visual perception is dampened, during the occurrence of an additional source of information (baroreceptors firing in systole). Conversely, they seem to gather somewhat more visual sensory information during the quiescent period of the cardiac cycle (in the absence of inner signalling and during the postsaccadic momentum). This idea is coherent with the presence of saccadic enhancement –increase vision before and after saccades (Ibbotson & Krekelberg, 2011), as well as to previous studies reporting dampened detection of pain, visual and auditory stimuli during systole compared to an improved perceptual sensitivity in diastole (Edwards et al., 2001, 2007; McIntyre et al., 2006; Pramme et al., 2014).

Overall, this work aimed to examine the phase-relationship between generated oculomotor events and the cardiac cycle (i.e., consistency of these events along the cardiac cycle). We found very consistent results across three different analyses (circular statistics, ANOVA and inter-trial-phase-coherence in Supp. materials). Although a consistent phase-relationship effect was found at the group level, it should be highlighted that this effect is driven by a very small proportion of the total number of oculomotor events (see ANOVA and subsequent table, as well as inter-trial-phase-coherence analysis in Supp. materials). Despite the effect size in terms of number of oculomotor events is not large, these are in the order of magnitude of previous studies examining changes in participants’ responses to stimuli locked to different cardiac phases (i.e., percentage change across responses; e.g., Garfinkel et al., 2013, 2014; Pramme et al., 2016). Another study also indicated a link between eye movements and heartbeat. Ohl et al., 2016 reported the coupling of microsaccades (small involuntary eye movements; ~< 1.5 °) with the systolic phase. The authors suggested that microsaccades may be modulated by baroreceptors via afferent signalling to the superior colliculus. In addition, as they also suggested a more parsimonious explanation: cardioballistic fluctuations of the ejected blood may exert a direct effect in the muscle activity (Fallon, 2004). Interestingly, the afferent discharge of muscle spindles is modulated by arterial pulsations (Birznieks, Boonstra, & Macefield, 2012)

## 6. Conclusions

By coregistration of eye movements and ECG, we found statistical evidence for a significant coupling of oculomotor behaviour with the heartbeat. Our results highlight the role of phasic changes within the cardiac cycle, and potentially, the cardioballistic ejection of the blood across the body, in modulating behaviour. More saccades were generated during the systolic phase and the concurrent upstroke of cardiac signalling. Contrary to this, fixations were more phase-locked to the quiescent period of the cardiac cycle. This is congruent with the hypothesis of humans sampling the external environment during quiescent periods of inner bodily signals such as heartbeat. We believe that these results provide original evidence for a mechanism by which humans might sample the world according to our bodily signals (i.e., a signal-to-noise ration account).

## Acknowledgments

We would like to thank all those who participated in and helped to advance this study. This research was supported by the Leverhulme Trust – Grant code RPG-2016-120

**Alejandro Galvez-Pol**: Conceptualization, Investigation, Methodology, Software, Formal analysis, Visualization, Writing original draft, Writing-Reviewing and Editing **R. McConnell**: Investigation, Validation, Visualization **J.M. Kilner:** Conceptualization, Methodology, Software, Formal analysis, Visualization, Writing-Reviewing and Editing, Project administrator, Funding acquisition

## References

Azevedo, R., Badoud, D., & Tsakiris, M. (2017). Afferent cardiac signals modulate attentional engagement to low spatial frequency fearful faces. Cortex, 1–9. https://doi.org/10.1016/j.cortex.2017.06.016

Azevedo, R. T., Garfinkel, S. N., Critchley, H. D., & Tsakiris, M. (2017). Cardiac afferent activity modulates the expression of racial stereotypes. Nature Communications, 8, 13854. https://doi.org/10.1038/ncomms13854

Birznieks, I., Boonstra, T. W., & Macefield, V. G. (2012). Modulation of human muscle spindle discharge by arterial pulsations - functional effects and consequences. PLoS ONE, 7(4). https://doi.org/10.1371/journal.pone.0035091

Brainard, D. H. (1997). The Psychophysics Toolbox. Spatial Vision, 10(4), 433–436. https://doi.org/10.1163/156856897X00357

Burr, D. C., Morrone, M. C., & Ross, J. (1994). Selective suppresion of the magnocellular visual pathway during saccadic eye movements. Nature, 371, 511–513. https://doi.org/10.1038/371511a0

Cheng, A., Eysel, U. T., & Vidyasagar, T. R. (2004). The role of the magnocellular pathway in serial deployment of visual attention. European Journal of Neuroscience, 20(8), 2188–2192. https://doi.org/10.1111/j.1460-9568.2004.03675.x

Cornelissen, F. W., Peters, E. M., & Palmer, J. (2002). The Eyelink Toolbox: Eye tracking with MATLAB and the Psychophysics Toolbox. Behavior Research Methods, Instruments, and Computers, 34(4), 613–617. https://doi.org/10.3758/BF03195489

Craig, A. D. (2002). How do you feel? Interoception: the sense of the physiological condition of the body. Nature Reviews Neuroscience, 3(8), 655–666. https://doi.org/https://doi.org/10.1038/nrn894

Critchley, H. D., & Garfinkel, S. N. (2018). The influence of physiological signals on cognition. Current Opinion in Behavioral Sciences, 19, 13–18. https://doi.org/10.1016/j.cobeha.2017.08.014

Critchley, H. D., & Harrison, N. A. (2013). Visceral Influences on Brain and Behavior. Neuron, 77(4), 624–638. https://doi.org/10.1016/j.neuron.2013.02.008

Edwards, L., Ring, C., McIntyre, D., & Carroll, D. (2001). Modulation of the human nociceptive flexion reflex across the cardiac cycle. Psychophysiology, 38(4), 712–718. https://doi.org/10.1017/S0048577201001202

Edwards, L., Ring, C., McIntyre, D., Carroll, D., & Martin, U. (2007). Psychomotor speed in hypertension: Effects of reaction time components, stimulus modality, and phase of the cardiac cycle. Psychophysiology, 44(3), 459–468. https://doi.org/10.1111/j.1469-8986.2007.00521.x

Fallon, J. B. (2004). Stochastic Resonance in Muscle Receptors. Journal of Neurophysiology, 91(6), 2429–2436. https://doi.org/10.1152/jn.00928.2003

Friston, K., Adams, R. A., Perrinet, L., & Breakspear, M. (2012). Perceptions as hypotheses: Saccades as experiments. Frontiers in Psychology, 3(MAY), 1–20. https://doi.org/10.3389/fpsyg.2012.00151

Friston, K., Kiebel, S., Barlow, H. B., Feynman, R. P., Neal, R. M., Hinton, G. E., & Neisser, U. (2009). Predictive coding under the free-energy principle. Philosophical Transactions of the Royal Society of London. Series B, Biological Sciences, 364(1521), 1211–1221. https://doi.org/10.1098/rstb.2008.0300

Garfinkel, S N, Minati, L., Gray, M. A., Seth, A. K., Dolan, R. J., & Critchley, H. D. (2014). Fear from the heart: sensitivity to fear stimuli depends on individual heartbeats. The Journal of Neuroscience, 34(19), 6573–6582. https://doi.org/10.1523/JNEUROSCI.3507-13.2014

Garfinkel, Sarah N., Barrett, A. B., Minati, L., Dolan, R. J., Seth, A. K., & Critchley, H. D. (2013). What the heart forgets: Cardiac timing influences memory for words and is modulated by metacognition and interoceptive sensitivity. Psychophysiology, 50(6), 505–512. https://doi.org/10.1111/psyp.12039

Garfinkel, Sarah N., & Critchley, H. D. (2016). Threat and the Body: How the Heart Supports Fear Processing. Trends in Cognitive Sciences, 20(1), 34–46. https://doi.org/10.1016/j.tics.2015.10.005

Hamker, F. H., Zirnsak, M., Calow, D., & Lappe, M. (2008). The peri-saccadic perception of objects and space. PLoS Computational Biology, 4(2). https://doi.org/10.1371/journal.pcbi.0040031

Ibbotson, M., & Krekelberg, B. (2011). Visual perception and saccadic eye movements. Current Opinion in Neurobiology, 21(4), 553–558. https://doi.org/10.1016/j.conb.2011.05.012

Khalsa, S. S., Adolphs, R., Cameron, O. G., Critchley, H. D., Davenport, P. W., Feinstein, J. S., … Paulus, M. P. (2018). Interoception and Mental Health: A Roadmap. Biological Psychiatry: Cognitive Neuroscience and Neuroimaging, 3(6), 501–513. https://doi.org/10.1016/j.bpsc.2017.12.004

Khalsa, S. S., & Lapidus, R. C. (2016). Can interoception improve the pragmatic search for biomarkers in psychiatry? Frontiers in Psychiatry, 7, 121. https://doi.org/10.3389/fpsyt.2016.00121

Kleiner, M., Brainard, D. H., Pelli, D. G., Broussard, C., Wolf, T., & Niehorster, D. (2007). What’s new in Psychtoolbox-3? Perception, 36, S14. https://doi.org/10.1068/v070821

Koch, E. (1932). Die irradiation der pressoreceptorischen kleislaufreflexe, (11), 225–227.

Kreindler, A. (1946). Recherches experimentales sur les relations entre le sinus carotidien et le systeme nerveux central. Bulletin de La Section Scientifique Academie Roumaine, (28), 448.

Kunzendorf, S., Klotzsche, F., Akbal, M., Villringer, A., Ohl, S., & Gaebler, M. (2019). Active information sampling varies across the cardiac cycle. Psychophysiology, (May 2018), 1–16. https://doi.org/10.1111/psyp.13322

McIntyre, D., Edwards, L., Ring, C., Parvin, B., & Carroll, D. (2006). Systolic inhibition of nociceptive responding is moderated by arousal. Psychophysiology, 43(3), 314–319. https://doi.org/10.1111/j.1469-8986.2006.00407.x

Ohl, S., Wohltat, C., Kliegl, R., Pollatos, O., & Engbert, R. (2016). Microsaccades Are Coupled to Heartbeat. Journal of Neuroscience, 36(4), 1237–1241. https://doi.org/10.1523/JNEUROSCI.2211-15.2016

Park, H.-D., Bernasconi, F., Salomon, R., Tallon-Baudry, C., Spinelli, L., Seeck, M., … Blanke, O. (2017). Neural Sources and Underlying Mechanisms of Neural Responses to Heartbeats, and their Role in Bodily Self-consciousness: An Intracranial EEG Study. Cerebral Cortex, (February 2018), 1–14. https://doi.org/10.1093/cercor/bhx136

Parr, T., & Friston, K. J. (2018). Active inference and the anatomy of oculomotion. Neuropsychologia, 111(October 2017), 334–343. https://doi.org/10.1016/j.neuropsychologia.2018.01.041

Pelli, D. G. (1997). The VideoToolbox software for visual psychophysics: Transforming numbers into movies. Spatial Vision. https://doi.org/10.1163/156856897X00366

Pickering, M. J., & Clark, A. (2014). Getting ahead: Forward models and their place in cognitive architecture. Trends in Cognitive Sciences, 18(9), 451–456. https://doi.org/10.1016/j.tics.2014.05.006

Pramme, L., Larra, M. F., Schächinger, H., & Frings, C. (2014). Cardiac cycle time effects on mask inhibition. Biological Psychology, 100(1), 115–121. https://doi.org/10.1016/j.biopsycho.2014.05.008

Pramme, L., Larra, M. F., Schächinger, H., & Frings, C. (2016). Cardiac cycle time effects on selection efficiency in vision. Psychophysiology, 53(11), 1702–1711. https://doi.org/10.1111/psyp.12728

Ross, J., Burr, D., & Morrone, C. (1996). Suppression of the magnocellular pathway during saccades. Behavioural Brain Research, 80(1-2), 1–8. https://doi.org/10.1016/0166-4328(96)00012-5

Salomon, R., Ronchi, R., Donz, J., Bello-Ruiz, J., Herbelin, B., Martet, R., … Blanke, O. (2016). The Insula Mediates Access to Awareness of Visual Stimuli Presented Synchronously to the Heartbeat. Journal of Neuroscience, 36(18), 5115–5127. https://doi.org/10.1523/JNEUROSCI.4262-15.2016

Seth, A. K. (2013). Interoceptive inference, emotion, and the embodied self. Trends in Cognitive Sciences, 17(11), 565–573. https://doi.org/10.1016/j.tics.2013.09.007

