## Supplementary Materials for "Active sampling in visual search is coupled to the cardiac cycle"

In addition to the circular statistics, we analysed the data in a repeated-measures ANOVA. To achieve this, the change in the proportion of each oculomotor event along the cardiac cycle was calculated in 6 bins that were equally spaced throughout the R-R interval (i.e., bin 1 refers to the start of the heartbeat and bin 6 to the end, early systole and late diastole, respectively). Given the number of post-hoc analyses (i.e., comparing bins across each type of event), we depict multiple comparisons in three subsequent tables (Table 1). Mauchly's W was computed to check for violations of the sphericity assumption and Greenhouse–Geisser adjustments to the degrees of freedom were applied when appropriate.

We first analysed differences between the types of oculomotor event in a repeated measures ANOVA (i.e., change in proportion of generated oculomotor event [Saccades, Fixations, Blinks] X Bins); this analysis yielded significant results ($F_{(3.4,105)} = 2.919$, $p = 0.032$). Secondly, we analysed each oculomotor separately (e.g., Saccades X Bins). For saccade onset there was a significant main effect of bin ($F_{(3.7,115)} = 2.910$, $p = 0.028$, Figure 1a supplementary). For fixations there was a significant main effect of bin ($F_{(4.3,133)} = 6.933$, $p = 0.000007$, Figure 1b supplementary). For blink onset there was also a significant main effect of bin (Saccades X Bin; $F_{(3.3,104)} = 2.833$, $p = 0.036$, Figure 1c supplementary).

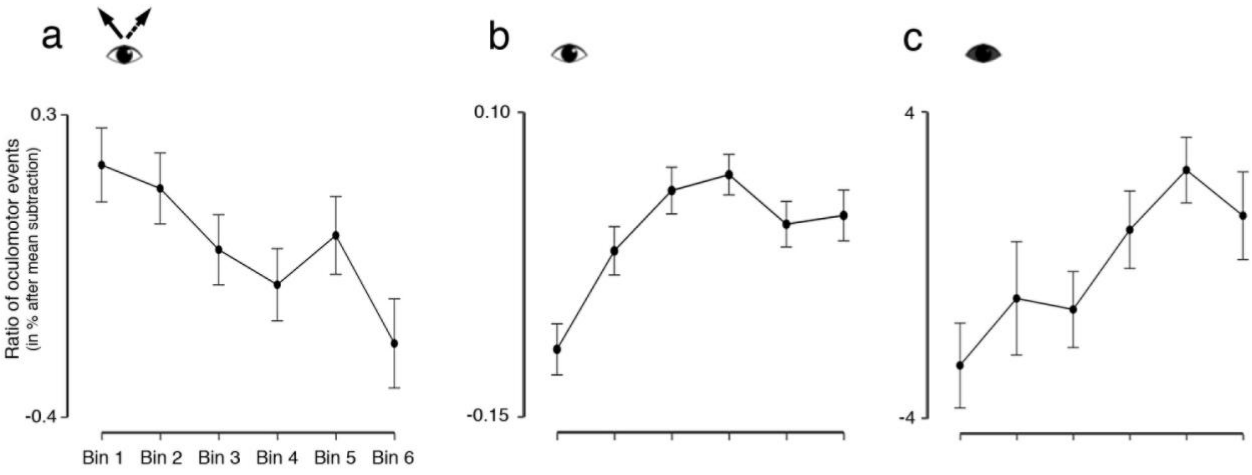


Figure 1 supplementary. Change in the proportion of Saccades, fixations, and blinks through the cardiac cycle. Panel A, B, C depict saccades, fixations and blinks, respectively. First, we calculated the proportion of oculomotor events that occurred in each bin (% of the total events across all bins). Next, to depict the change in percentage across bins, we subtracted the expected proportion if no change occurs (100% divided by 6 bins, null proportion) to the ratio of each bin; i.e., change in proportion of oculomotor events in % after mean subtraction. Error bars show the standard error of the mean; Bin 1 refers to the beginning of the heartbeat (early systole bounded by R-peak) and Bin 6 to the end of the heartbeat (late diastole bounded to next heartbeat R-peak).

Table 1.

*Post Hoc Tests by oculomotor event across 6 bins equally spaced throughout the R-R interval*

| | | Saccades | | | Fixations | | | Blinks | | |
|---|---|---|---|---|---|---|---|---|---|---|
| | | t | Cohen's d | p bonf | t | Cohen's d | p bonf | t | Cohen's d | p bonf |
| Bin 1 | Bin 2 | 0.426 | 0.075 | 1 | -2.74 | -0.484 | 0.151 | -0.878 | -0.155 | 1 |
| | Bin 3 | 1.895 | 0.335 | 1 | -5.144 | -0.909 | 2.134e -4 | -1.383 | -0.245 | 1 |
| | Bin 4 | 2.4 | 0.424 | 0.339 | -5.374 | -0.95 | 1.102e -4 | -2.092 | -0.37 | 0.671 |
| | Bin 5 | 1.636 | 0.289 | 1 | -3.519 | -0.622 | 0.02 | -3.422 | -0.605 | 0.026 |
| | Bin 6 | 2.604 | 0.46 | 0.21 | -3.444 | -0.609 | 0.025 | -2.644 | -0.467 | 0.191 |
| Bin 2 | Bin 3 | 1.136 | 0.201 | 1 | -1.605 | -0.284 | 1 | 0.142 | 0.025 | 1 |
| | Bin 4 | 1.949 | 0.345 | 0.906 | -2.569 | -0.454 | 0.228 | -1.107 | -0.196 | 1 |
| | Bin 5 | 0.928 | 0.164 | 1 | -0.794 | -0.14 | 1 | -2.306 | -0.408 | 0.419 |
| | Bin 6 | 3.061 | 0.541 | 0.068 | -1.108 | -0.196 | 1 | -1.031 | -0.182 | 1 |
| Bin 3 | Bin 4 | 0.632 | 0.112 | 1 | -0.478 | -0.084 | 1 | -1.368 | -0.242 | 1 |
| | Bin 5 | -0.285 | -0.05 | 1 | 1.224 | 0.216 | 1 | -2.652 | -0.469 | 0.187 |
| | Bin 6 | 1.749 | 0.309 | 1 | 0.688 | 0.122 | 1 | -1.84 | -0.325 | 1 |
| Bin 4 | Bin 5 | -0.86 | -0.152 | 1 | 1.607 | 0.284 | 1 | -1.331 | -0.235 | 1 |
| | Bin 6 | 1.183 | 0.209 | 1 | 1.38 | 0.244 | 1 | -0.25 | -0.044 | 1 |
| Bin 5 | Bin 6 | 1.605 | 0.284 | 1 | -0.243 | -0.043 | 1 | 0.79 | 0.14 | 1 |

*Note.* P values are corrected for multiple comparisons using Bonferroni correction (p bonf).

We also calculated the inter-trial-phase-coherence, whereby a vector of a certain length and direction represents the strength of the observed effects. To this aim, the phase of each event regarding the saccade and blink onset in the R-R cycle was calculated for each subject. For every event this produced values with a phase and a vector length set to 1. These polar coordinates were transformed into Cartesian coordinates and the average vector was calculated for each subject; the length of this vector corresponded to the strength of the effect (bounded by 0 and 1). If all events had the same phase then the vector length would be equal to 1 whereas if the events were evenly distributed around the circle then the length would be 0. To determine whether these vector lengths were significant we generated a simulated dataset. To achieve this, we randomly sampled from a uniform distribution of phases between 0 and 2π. The number of events sampled were the same as the number of events for the actual blink and saccade data and this was repeated 32 times. In this way, a null data set was created with the same number for events and participants as the actual data. For this data the inter-trial-phase-coherence was calculated in the same way as above. This simulation was run 10,000 times and the distribution of the null space was produced. The actual value was compared to this distribution as was considered significant if it was in the 2.5% tails, equivalent to $p<0.05$.

For blink onset, the mean inter-trial-coherence was 0.138 (std 0.11). This was significantly greater than the null distribution for blinks (Figure 2a supplementary). For saccade onset the mean inter-trial-coherence was 0.007 (std 0.003). This was significantly greater than the null distribution for saccades (Figure 2b supplementary). These data corroborate the previous phase analysis and the ANOVA, confirming both a significant phase relationship between saccades, blinks, and the cardiac cycle, as well as confirming that although consistent, this is a relatively small effect size in terms of number of saccades and blinks.

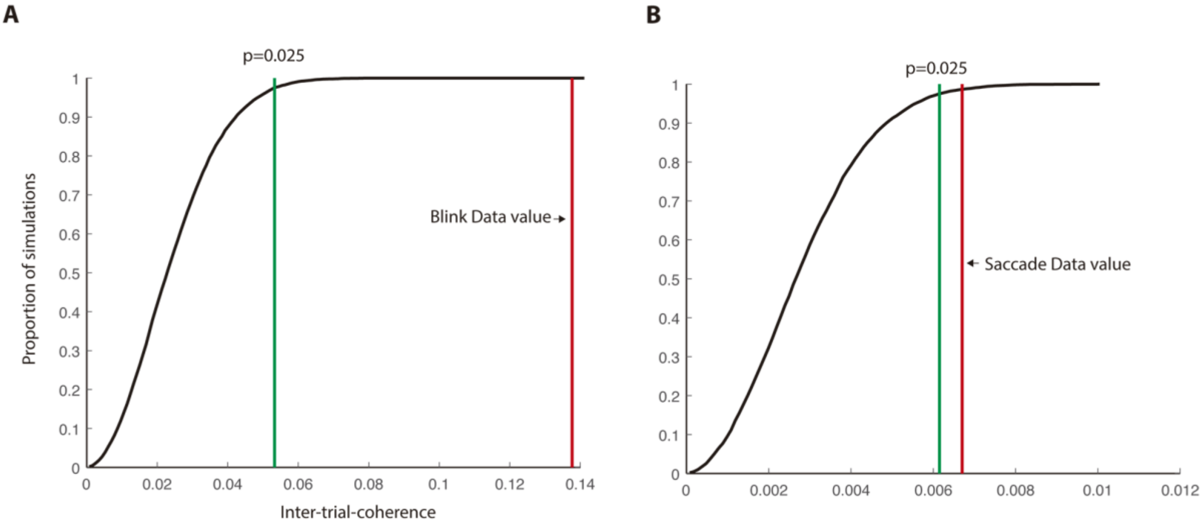


Figure 2 supplementary. Inter-trial-phase-coherence; panel A and B refer to the blink and saccade dataset, respectively. For blink onset, the mean inter-trial-coherence was 0.138 (std 0.11), which was significantly greater than the null distribution for blinks. For saccade onset the mean inter-trial-coherence was 0.007 (std 0.003). This was significantly greater than the null distribution for saccades. Green line bounds with the null distribution whereby the red line depicts the experiment's dataset.
